# A refinement to gene editing in Atlantic salmon using asymmetrical oligonucleotide donors

**DOI:** 10.1101/2021.02.08.430296

**Authors:** Anne Hege Straume, Erik Kjærner-Semb, Kai Ove Skaftnesmo, Hilal Güralp, Simon Lillico, Anna Wargelius, Rolf Brudvik Edvardsen

## Abstract

Selective breeding programs in aquaculture are limited by the heritability of the trait and long generation time in most fish species. New breeding technology (NBT) using CRISPR/Cas9-induced homology directed repair (HDR) have the potential to expedite genetic improvement in aquaculture, but the method requires optimization. Here we show that asymmetrical oligonucleotide (ODN) donors induce highly efficient and precise edits in individual Atlantic salmon founder animals. We performed single nucleotide replacement (SNR) in *dnd* with up to 59.2% efficiency, and inserted FLAG elements into *slc45a2* and *dnd*, with up to 36.7 % and 32.7% efficiency, respectively. We found HDR efficiency to be dependent on template concentration, but a trade-off with respect to toxicity was observed. Using this NBT in salmon we demonstrate that precise modification of the genome can be achieved in a single generation, allowing efficient introgression of favorable alleles and bypassing challenges associated with traditional selective breeding.

## Introduction

There is an increasing demand for sustainable animal husbandry, and the fast-growing fish aquaculture industry is a food production sector with great potential to improve global food security. Fish aquaculture is also considered to be efficient in terms of feed conversion and protein retention compared to most terrestrial livestock^1,2^. Atlantic salmon (*Salmo salar* L.) is farmed in the sea at a large scale, but further growth is currently hindered by a range of issues including genetic introgression of escapees into wild populations and the spread of disease^3,4^. New breeding technology (NBT) using gene editing offer an exciting opportunity to increase the sustainability of open sea-cage salmon farming by allowing us to induce both sterility and disease resistance^2,5-7^. An important issue when it comes to gene editing in salmon, is to reduce mosaicism in the founder fish. The long generation time (3-4 years) makes breeding an unattractive option to obtain homozygous mutants, and most functional studies must be performed in F0. However, to produce homozygous F1 fish by intercrossing, it will also be desirable to obtain a high percentage of perfect editing in individual F0 fish. Thus, improving the efficiency of precise editing in founder individuals is more important than obtaining a high number of mosaic F0 fish. A CRISPR/Cas9 induced double-stranded DNA break (DSB) in the coding sequence of a gene, followed by activation of the endogenous non-homologous end joining (NHEJ) pathway, results in an array of unpredictable insertions or deletions that may result in frameshift and gene knock-out (KO). This is a useful approach to study KO phenotypes, and has been applied successfully in salmon^5,7,8^ and several other farmed fish species^9-24^. To make precise genome alterations it is a necessity to induce homology directed repair (HDR) by supplying a repair template with homology to the CRISPR target site, thereby allowing to change SNPs, insert affinity tags for protein detection and modify regulatory elements to alter expression of target genes. A single nucleotide replacement (SNR) can be used to introduce favorable naturally alleles and could be a promising and time saving solution compared to traditional breeding with backcrossing and selection. The genetic progress in selective breeding programs is also limited by the heritability of the target traits, and the standing genetic variation in the broodstock. NBT using CRISPR/Cas9-induced HDR can offer new solutions and opportunities in these areas^2,25^.

We have previously demonstrated highly efficient HDR in salmon using symmetrical oligonucleotides (ODNs) with short (24/48/84 bp) homology arms to knock-in (KI) a FLAG element in the pigmentation gene *solute carrier family 45 member 2* (*slc45a2)*. Using high-throughput sequencing (HTS), we showed *in vivo* ODN-mediated KI in almost all the gene edited animals and demonstrated perfect HDR integration rates of up to 27 % in individual F0 embryos^26^. Short homology arms have also been shown to induce efficient HDR in zebrafish^27,28^.

In this work we aimed to further improve the HDR precision and efficiency in salmon, with the goal to reduce mosaicism in individual F0 animals. Asymmetrical ODNs in combination with CRISPR/Cas9 have previously been demonstrated to improve HDR rates in cell cultures^29^ and induced pluripotent stem cells^30^. Based on these promising results, we have explored the use of asymmetrical ODNs. We have successfully performed a SNR in the primordial germ cell survival factor gene *dead end* (*dnd*), and inserted FLAG elements into both *slc45a2* and *dnd*. SNR was more efficient than FLAG KI, suggesting that HDR efficiency may be inversely proportional with insert size. As previously^26^, we found HDR efficiency to be dependent on template concentration, but suggest using the lowest possible concentration to avoid toxicity and enable targeting multiple genes at the same time. Our results show that CRISPR/Cas9 in combination with asymmetrical ODNs enables rapid and precise changes to the genome in individual F0 animals and present a promising tool for fish breeders in the future.

## Results and Discussion

### FLAG KI targeting slc45a2 and dnd

Targeting *slc45a2*^*5*^and *dnd*^*7*^, we have here performed KI of a FLAG element in F0 salmon using CRISPR/Cas9 and asymmetrical ODNs (Fig. 1a and Supplementary Fig. 1). Analyzing the percentage of perfect HDR in individual animals by HTS of amplicons, we detected an average of 13.6 % (std 10.9 %) for *slc45a2* and 7.6 % (std 10.1 %) for *dnd* (Fig. 1b). Interestingly, we observed some individuals with a very high efficiency in both groups with up to 36.7 % perfect HDR in *slc45a2*, and 32.7 % in *dnd* (Supplementary Table 1). This is higher than our previously reported results showing an average of up to 6.7 % perfect FLAG KI in *slc45a2*, and a maximum of up to 26 % perfect HDR in individual animals, using symmetrical ODNs at 1.5 µM^26^. Comparing the efficiency of FLAG KI (targeting *slc45a2*) using asymmetrical ODNs described herein, to symmetrical ODNs described before^26^, a significant difference was detected for average perfect HDR between symmetrical (5.1 %) and asymmetrical ODNs (13.6 %). No significant difference was detected when comparing he average rates of erroneous HDR between symmetrical (3.1 %) and asymmetrical ODNs (2.0 %). (Supplementary Fig. 2).

**Fig 1:**
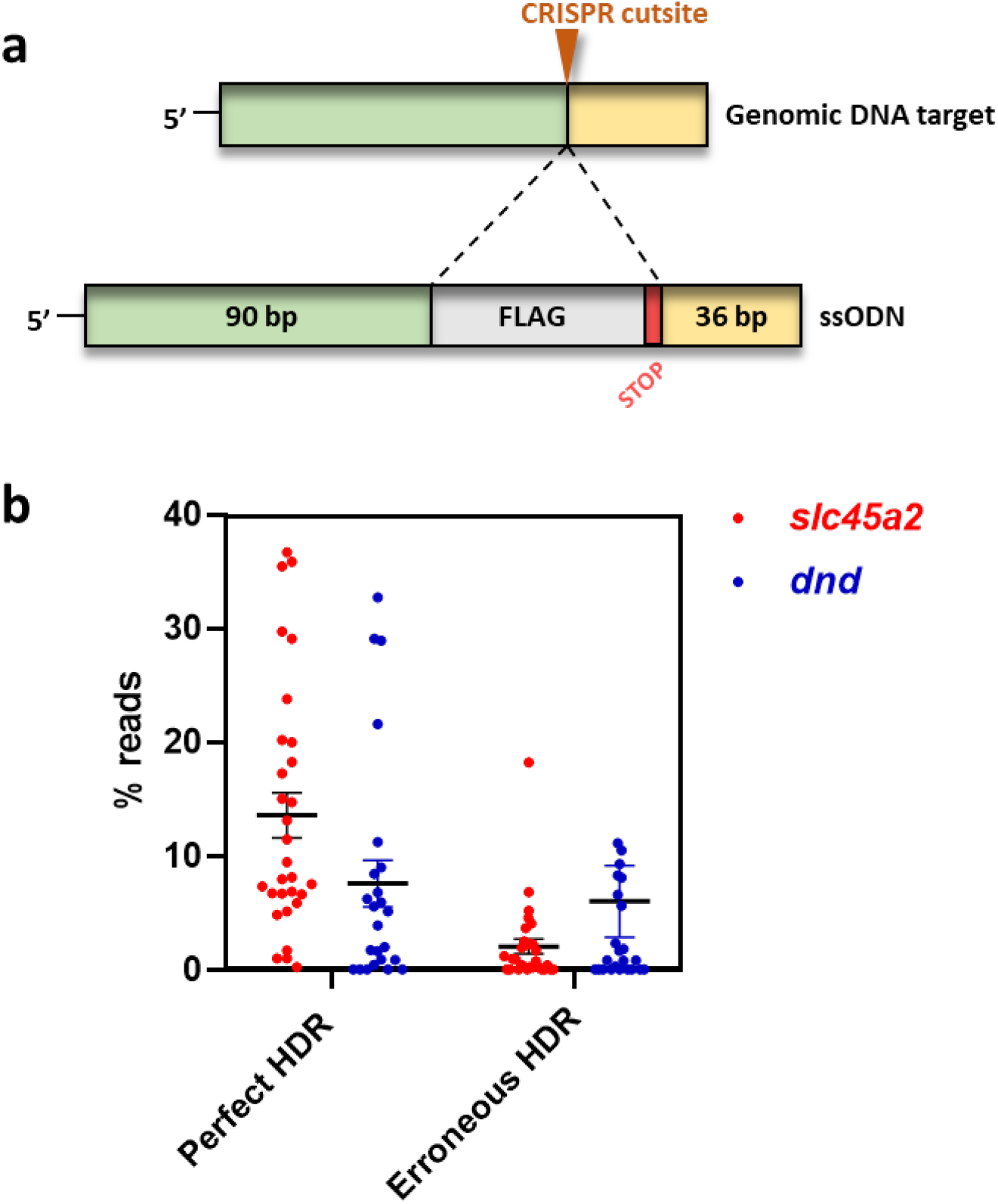
*slc45a2* and *dnd* FLAG knock-in (1.5 µM ODN). **a** Asymmetrical ODNs were designed by copying 90 + 36 nucleotides on each side of the CRISPR cut site flanking the insert (indicated with a dotted line) containing the FLAG element followed by a STOP codon (TAA). **b** Relative read counts per individual for *slc45a2* (red dots, n=30) and *dnd* (blue dots, n=24). Reads with a perfect match to the entire target sequence are referred to as perfect HDR. Reads with a correct insert flanked by mismatches/indels on the 5’ and/or 3’-side are referred to as erroneous HDR. Error bars indicate SEM/group.

### Oligonucleotide concentration

We and others^26,31^ have shown that increasing the concentration of the DNA donor improves HDR efficiency. However, DNA can be toxic to cells and we wanted to elucidate if there is a trade-off between high integration efficiency and toxicity by testing the *slc45a2* FLAG KI ODN at three different concentrations: 0.5, 1.5 and 4.0 µM (Fig. 2). In accordance with our previous results, a template concentration of 1.5 µM resulted in the most efficient KI. We detected the approximately same average efficiency when using 0.5 and 4 µM. However, the highest concentration resulted in fewer pure albinos and a higher degree of mosaicism compared to individuals injected with lower concentrations of template (Supplementary Fig. 3). As expected, the HTS results from the animals who had received the highest dose revealed a much higher percentage of wild type reads (Supplementary Table 1). During the microinjection procedure there will be inevitable variation in the volume injected into each fertilized egg. Performing precise microinjections by hand can be challenging due to the opaque salmon eggs, and personal skills will influence the outcome. Technical aspects will also matter, such as variation in the diameter of the needle opening and the egg quality. It is therefore conceivable that the mosaicism observed for the high dose group (4 µM ODN) is due to toxicity of the injection mix when the injected volume is high. We hypothesize that the surviving eggs received a lower volume of the injection mix, but as they also received a lower dose of the *cas9*- and guide RNA they became more mosaic. Taking this into account, it could be an advantage to use the lowest possible ODN concentration to avoid unnecessary DNA induced toxicity, which would allow editing multiple genes at the same time.

**Fig 2:**
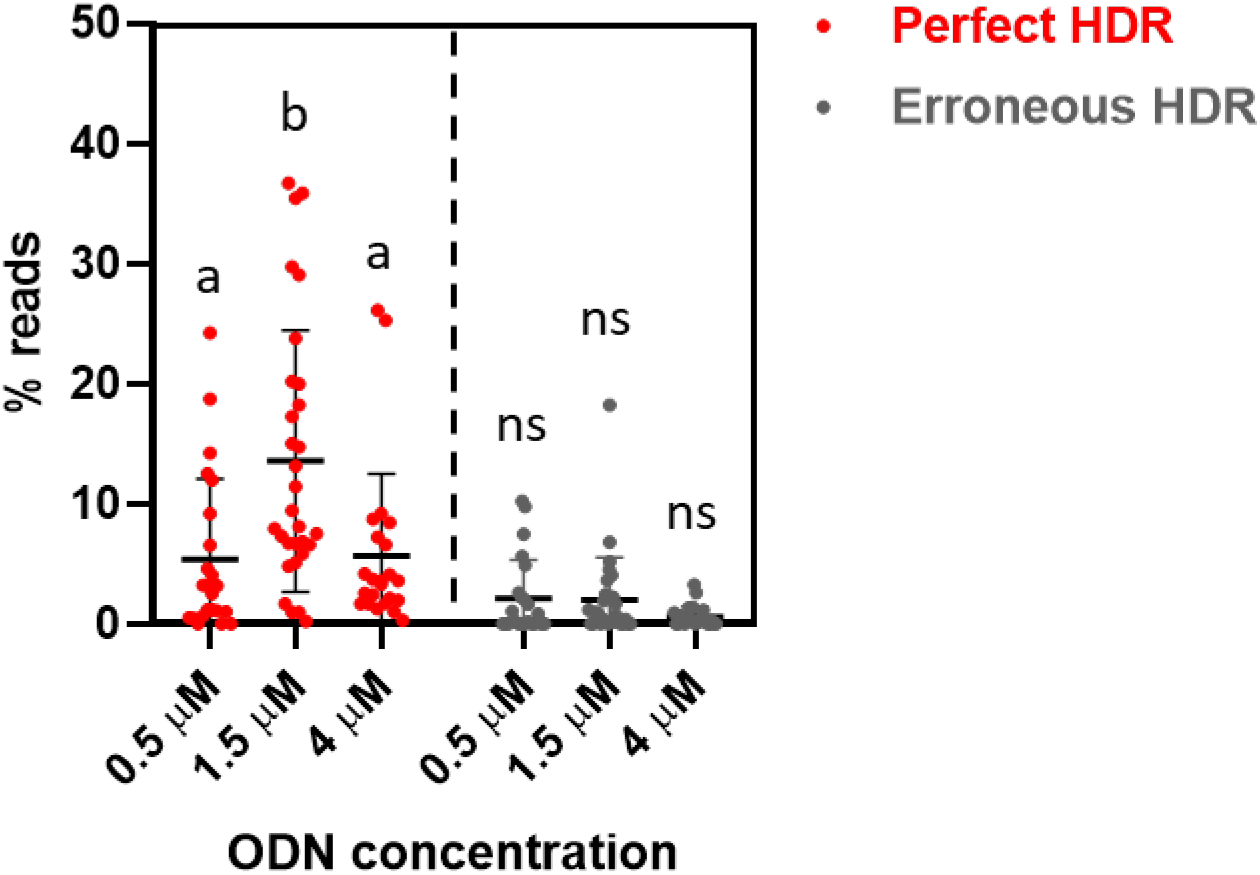
*slc45a2* FLAG knock-in. The asymmetrical ODN targeting *slc45a2* was tested using three different concentrations: 0.5 (n=23), 1.5 (n=30) and 4.0 (n=23) µM. Sequence reads with a perfect match to the entire target sequence are referred to as perfect HDR and reads with a correct insert but mismatches/indels in the homology arms are referred to as erroneous HDR. Read counts for each sample are given in % of the total number of reads. The error bars indicate SEM/group. Different lowercase letters indicate significant differences (*P* < 0.05), ns = non-significant.

### Single Nucleotide Replacement

Targeting *dnd*, we performed a SNR using an asymmetrical ODN while at the same time continuing to refine the ODN concentration. Using 0.15, 1.5 and 4 µM ODN concentrations, we obtained an average perfect HDR of 7.4 % (std 14.8), 12.5 % (std 14.3) and 7.4 % (std 9.4), respectively (Fig. 3). However, when analyzing individual fish, the most striking result was obtained using 1.5 µM were we detected perfect repair efficiency up to 59.2 %. To our knowledge, this level of perfect HDR in F0 has not been reported in any other fish. Even at the lowest concentration (0.15 µM), two individuals displayed 49.1 and 47.4 % perfect HDR. We speculate that the high efficiency for SNR is due to the lack of insert, as editing efficiency has been shown to be sensitive to insert size^32^. When CRISPR/Cas9 is used to make a traditional KO through NHEJ, one of the challenges is that the mutation can be in-frame and therefore potentially silent. SNR could solve this by insertion of novel stop codons and as such increase the levels of functional KO mutations. Moreover, for some genes and applications, it may not be relevant to perform KI but to make smaller edits such as a changing one or a few SNPs. One such example is the *vgll3* locus containing two missense SNPs strongly linked to age at maturity in salmon^3^. Developing precise gene editing technology to make such small edits may therefore be useful to enable NBT introgression of natural beneficial variants into aquaculture strains.

**Fig 3:**
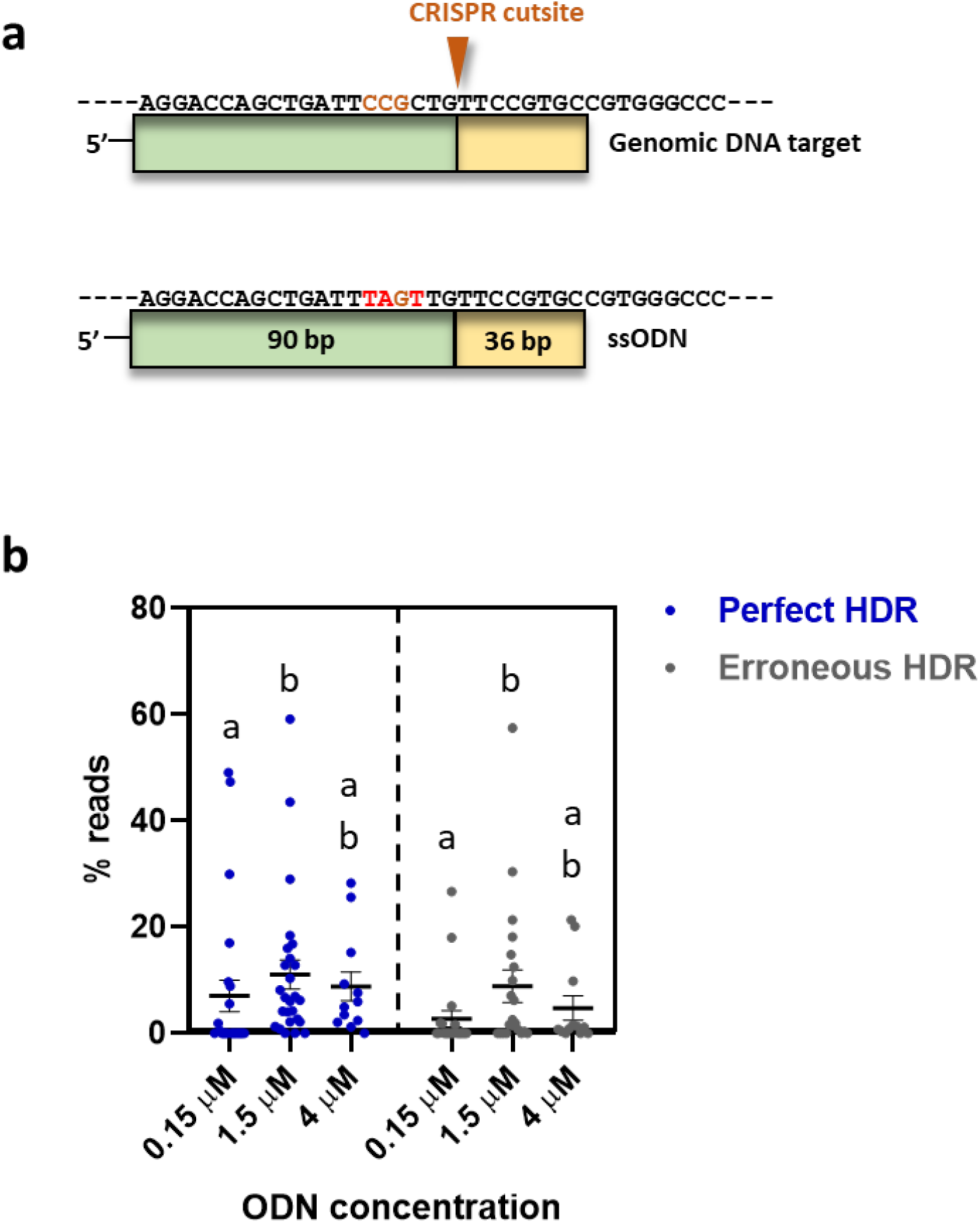
Single nucleotide replacement in *dnd*. **a** An asymmetrical ODN targeting *dnd* was designed with 90 + 36 nucleotides on each side of the CRISPR cut site and three nucleotides were changed. PAM site is shown with brown letters, and novel nucleotides with red letters. **b** HDR rates for three different ODN concentrations; 0.15 (n=24), 1.5 (n=26) and 4.0 µM (n=12). Sequence reads with a perfect match to the entire target sequence are referred to as perfect HDR (blue) and reads with a correct SNR but mismatches/indels in the homology arms are referred to as erroneous HDR (gray). Read counts for each sample are given in % of the total number of reads. Error bars indicate SEM/group. Different lowercase letters indicate significant differences (*P* < 0.05).

### Erroneous repair and Indel locations

In addition to reads displaying perfect HDR, we detected reads displaying erroneous repair, meaning reads with a correct FLAG-insert/SNR but also indels on the 5’- and/or 3’-side of the insert/SNR (Figs. 1b, 2 and 3b).

In our previous work using symmetrical ODNs, we revealed a strong correlation between ODN polarity and the location of these indels on either the 5’- or 3’-side of the inserted sequence^26^. According to this, most of the indels will end up on the 5’-side of the insert, when using a repair template with sense orientation relative to the target strand (reverse complementary to the gRNA) (Supplementary Fig. 1a and b). In the current study the orientation of the asymmetrical ODNs were sense relative to the target strand, and we observed that most of the indels were indeed located on the 5’-side of the insert (fewer reads with perfect 5’-reads than 3’-reads). Although this finding supports our previous results, we only detected a significant difference for *dnd* KI and SNR in the present study (Fig. 4).

**Fig 4:**
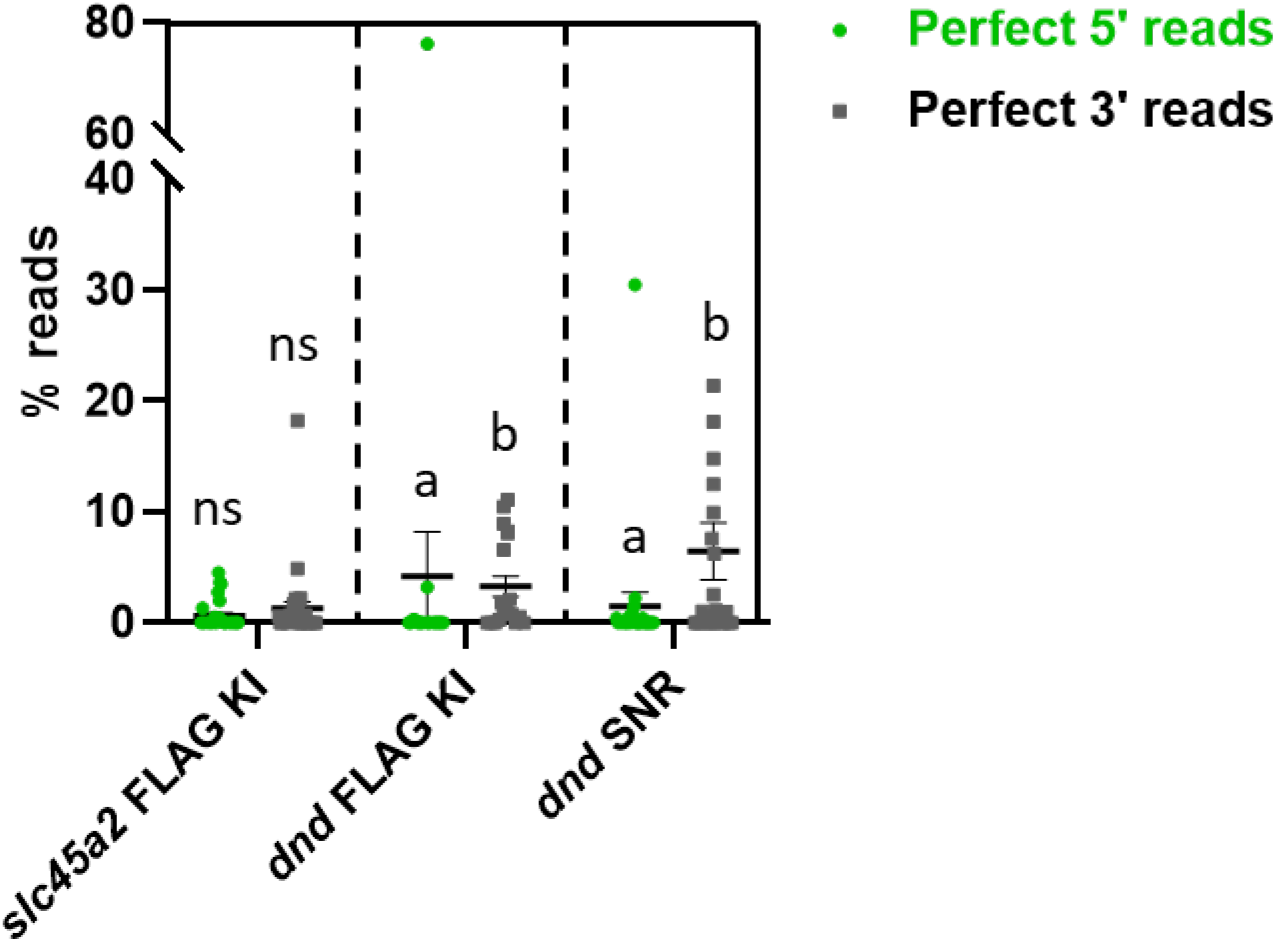
Variation in indel locations. Here, we distinguished between reads with a perfect match to the 5’- <u>or</u> 3’-side of the FLAG insert/SNR. The asymmetrical ODNs were compared at 1.5 µM. Green dots represent perfect 5’ reads and squares represent perfect 3’ reads. Read counts for each sample are given in % of the total number of reads. Error bars indicate SEM/group. The groups *slc45a2* FLAG KI (n = 30), *dnd* FLAG KI (n = 19) and *dnd* SNR (n = 24) were analyzed separately. Different lowercase letters indicate significant differences (*P* < 0.05).

We have here demonstrated that asymmetrical ODNs induce efficient and precise HDR in salmon, both for KI- and SNR. Moreover, asymmetrical ODNs appear to be more efficient than symmetrical ones, as compared to our previous results^26^ (Supplementary Fig. 2). Comparing the outcome of SNR and KI in *dnd* in we found SNR to be the most efficient approach, suggesting that HDR efficiency is inversely proportional with insert size. Although we found HDR efficiency to be dependent on template concentration, it might be beneficial to use the lowest possible template concentration to avoid toxicity and enable editing multiple genes at the same time. We show that it is possible to use CRISPR/Cas9-induced HDR in NBT to obtain desirable traits. SNR is a promising tool to insert favorable alleles in farmed salmon and, considering the long generation time, more convenient than crossing in traits through conventional breeding. Moreover, this could also be an advantage for aquaculture species in general (e.g trout, sea bass, tilapia). This technology offers an exciting opportunity to insert traits of interest into the recently demonstrated fertile but genetically sterile salmon^25^. This fish will produce sterile offspring and may therefore represent the future salmon aquaculture by combining sterility and other favorable traits induced by HDR, such as disease resistance.

## Methods

### Ethics statement

This experiment was approved by the Norwegian Animal Research Authority (NARA, permit number 14865) and the use of these experimental animals was in accordance with the Norwegian Animal Welfare Act.

### Preparation of Cas9 RNA, gRNAs and ODNs

The CRISPR target sequences for *slc45a2* and *dnd1* are described in Edvardsen et al.^5^, and Wargelius et al.^7^, respectively. Preparation of gRNAs and *cas9* mRNA was performed as previously described^5,26^. The RNeasy MiniKit spin column (Qiagen) was used to purify the gRNA. The ODNs were ordered from Integrated DNA Technologies (Leuven, Belgium). The ODN design is based on Richardson et al.^29^.

### Microinjection

Salmon eggs and sperm were delivered by Mowi (Hauglandshella, Askøy, Norway). Fertilization and microinjections were carried out as described previously^5^ using 50 ng/µl gRNA and 150 ng/µl *cas9* mRNA in nuclease free water and a FemtoJet®4i (Eppendorf) microinjector. The ODNs were added to the injection mix with a final concentration of 0.15, 0.5, 1.5 or 4 µM.

### Analysis of mutants

As described previously^26^ *slc45a2* mutants were selected based on visual inspection of newly hatched larvae. When editing *dnd*, we also added the *slc45a2* gRNA to the injection mix to obtain a visual phenotype, and thus make it easier to select the mutants. DNA was extracted from caudal fins using DNeasy Blood & Tissue kit (Qiagen). DNA extracted from the fin has previously been shown to be broadly representative for the whole fish^5,25^. A fragment covering the entire CRISPR target sites for *slc45a2* and *dnd1* was amplified with a two-step fusion PCR (as described in Gagnon et.al 2014) to prepare for Illumina MiSeq. The following primers (gene specific sequence indicated in capital letters) were used in the first PCR-step for *slc45a2*: 5’-tctttccctacacgacgctcttccgatctCAGATGTCCAGAGGCTGCTGCT and 5’-tggagttcagacgtgtgctcttccgatctTGCCACAGCCTCAGAATGTACA. The following primers (gene specific sequence indicated in capital letters) were used in the first PCR-step for *dnd*: 5’-tctttccctacacgacgctcttccgatctGGGGAAAGGCTAGGGAGAGA and 5’-tggagttcagacgtgtgctcttccgatct CGGTTCTGTCCGCTGAAGTT.

### Analysis of MiSeq data

Read counts were reported for variants containing the inserted or edited sequence, separating those with a perfect match to the entire target sequence (referred to as perfect HDR), and those with a correct insert sequence/SE, but mismatches in the rest of the target sequence (referred to as erroneous HDR). In addition, read counts were reported for wild type sequences. The settings applied for filtering, trimming and variant calling of the MiSeq reads are illustrated in Supplementary Fig. 4, and described below:

Fastq files were filtered and trimmed with the following specifications; primer sequences were used to demultiplex reads from different amplicons on the same sequencing run, minimum read length was set to 100 bp, and forward and reverse reads were assembled to correct sequencing errors (minimum overlap between forward and reverse reads was set to 150 bp for *slc45a2* and 200 bp for *dnd*, and allowing maximum 20% mismatches between forward and reverse reads in the overlap region). Assembled reads were combined with forward reads that did not pass the assembly thresholds. Variants were then called using positions 20-200 for *slc45a2* and positions 60-230 for *dnd*. All bases with base quality < 20 were converted to N’s, and maximum 5 N’s were allowed per read. Identical reads were then grouped (referred to as variants), and variants that only differed by up to 5 N’s were grouped if none of the variants differed by any nucleotides. For each group, the variant with the least N’s was chosen as representative. We only retained variants supported by a minimum of 100 reads and variants were grouped if they differed by up to 5 N’s if none of the variants differed by any nucleotides.

### Statistical analyses

D’Agostino Person normality test (column statistics) were used to assess normal distribution of the data. None of the groups displayed normal distribution, and we carried on with non-parametric analyses. When analyzing more than two groups, non-parametric statistical analyses were performed using a Kruskall-Wallis test, followed by Dunn’s multiple comparison test. When analyzing two groups, a Mann-Whitney rank test, or a Wilcoxon paired test was performed. The tests were carried out using GraphPad Prism 8.0.1.

## Acknowledgement

Our study was funded by the EU-COFASP project: “AQUACRISPR: Optimization of the CRISPR/Cas9 knock-in technology and application in salmon and trout”. We would like to thank Lise Dyrhovden and Ivar Helge Matre for their assistance in rearing. We are also grateful for the assistance of Hanne Sannæs and Ida Kristin Mellerud for running the MiSeq, Aquagen (Trondheim, Norway) and Mowi (Hauglandshella, Askøy, Norway) for providing eggs and sperm.

## Author contributions

R.B.E., A.H.S. and A.W. designed the project. A.H.S., E.K.S., H.G. and R.B.E. performed the microinjections. A.H.S. and R.B.E. designed the ODN templates and collected the tissue samples. A.H.S. made the gRNA and Cas9 RNA, purified the DNA and ran the PCR screening. K.O.S. prepared the Illumina sequencing libraries. E.K.S. performed the bioinformatic analysis of the NGS. A.H.S., A.W. and R.B.E. analyzed the results and wrote the paper. All authors read and approved the final manuscript.

## Additional Information

### Data Availability

Data generated or analyzed during this study are included in this article (and its Supplementary Information).

