## Supplementary figures for "A refinement to gene editing in Atlantic salmon using asymmetrical oligonucleotide donors"

Supplementary Figures 1-4

**Supplementary Fig. 1:**

**a** Asymmetrical ODN design *slc45a2*

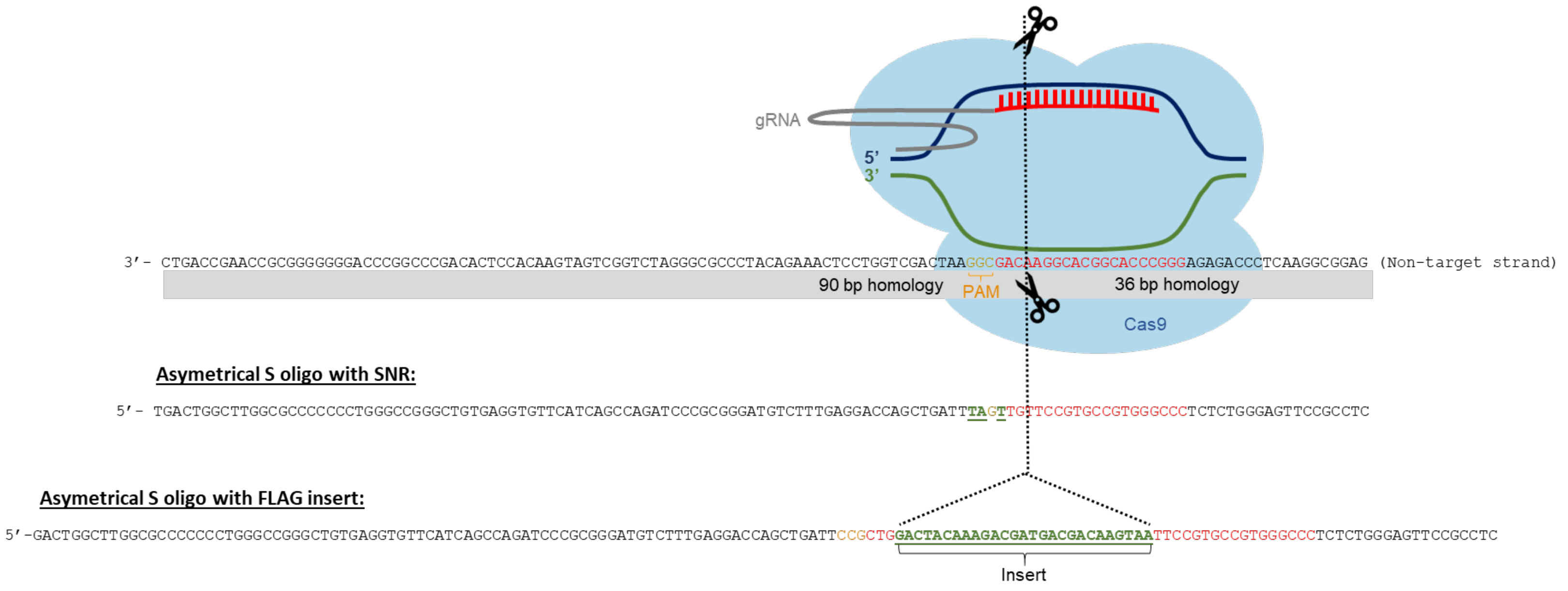


**b** Asymmetrical ODN design *dnd*

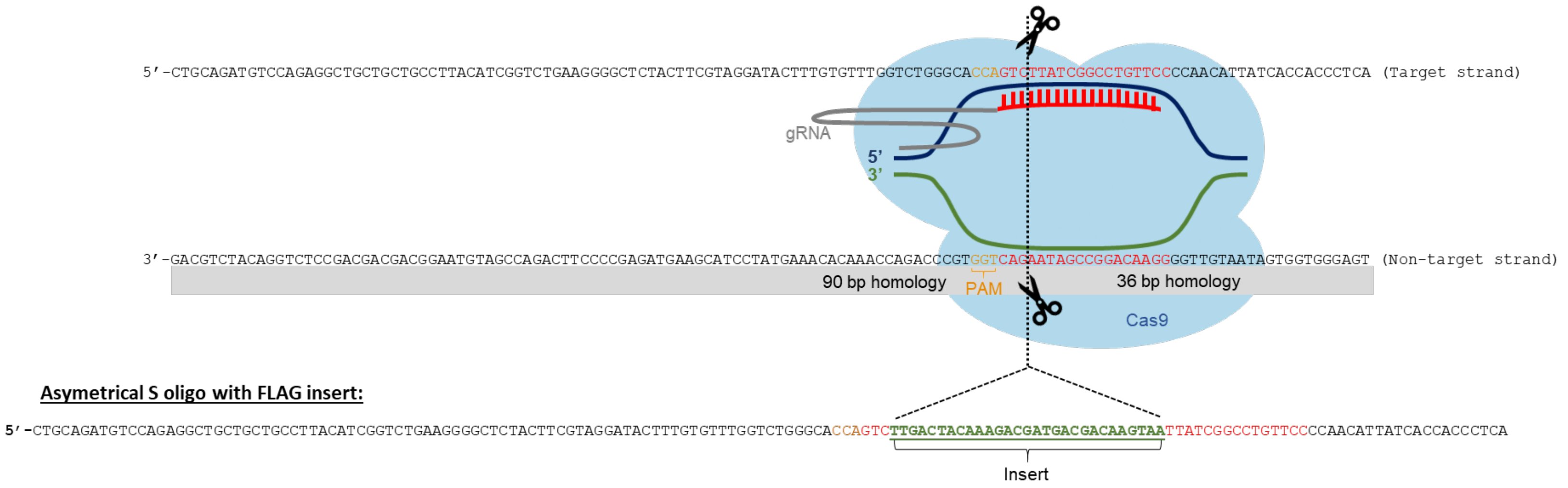

**Supplementary Fig. 2:**

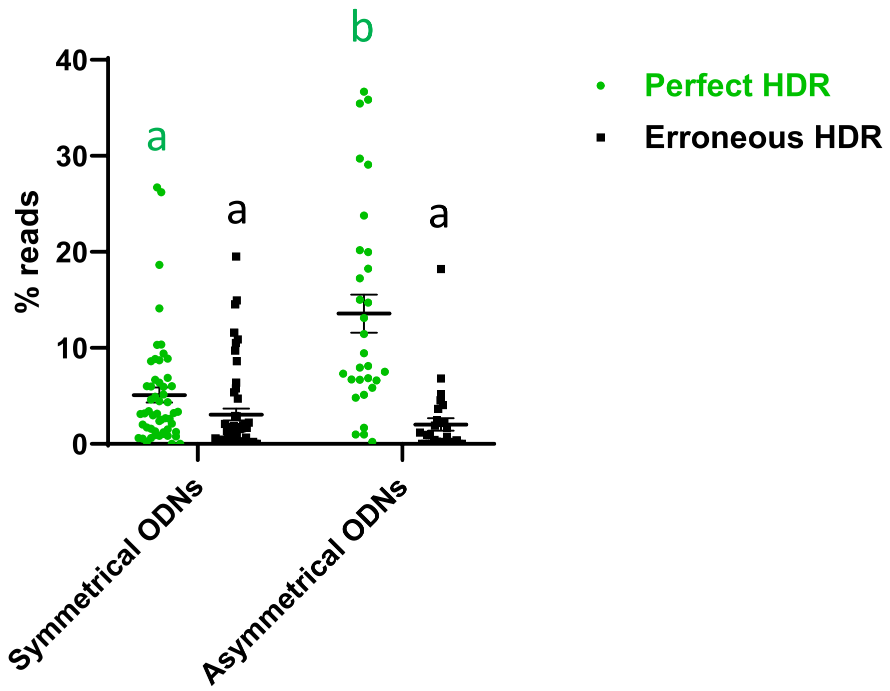


**Comparison of symmetrical and asymmetrical ODNs for *slc45a2* FLAG knock-in**

All the ODNs compared here were designed for *slc45a2* to KI a FLAG element. The ODN concentration was 1.5 µM. The symmetrical ODNs are a pool of S 24, AS 24, ds 24, S 48 and S 84 (described in Straume et.al 2020). The asymmetrical ODN design is illustrated in Suplemmentary Fig. 1 A. Green dots represent perfect HDR, black squares represent erroneous HDR. Mutant fish were analysed using Illumina MiSeq. Read counts for each sample are given in % of the total number of reads with at least 100 identical reads. The error bars indicate the SEM of the mean for each group. A Mann-Whitney test was used to compare the mean rank of symmetrical vs. asymmetrical ODNs, analyzing the groups perfect and erroneous HDR separately. Different lower-case letters indicate significant differences ($P < 0.05$).

**Supplementary Fig. 3.**

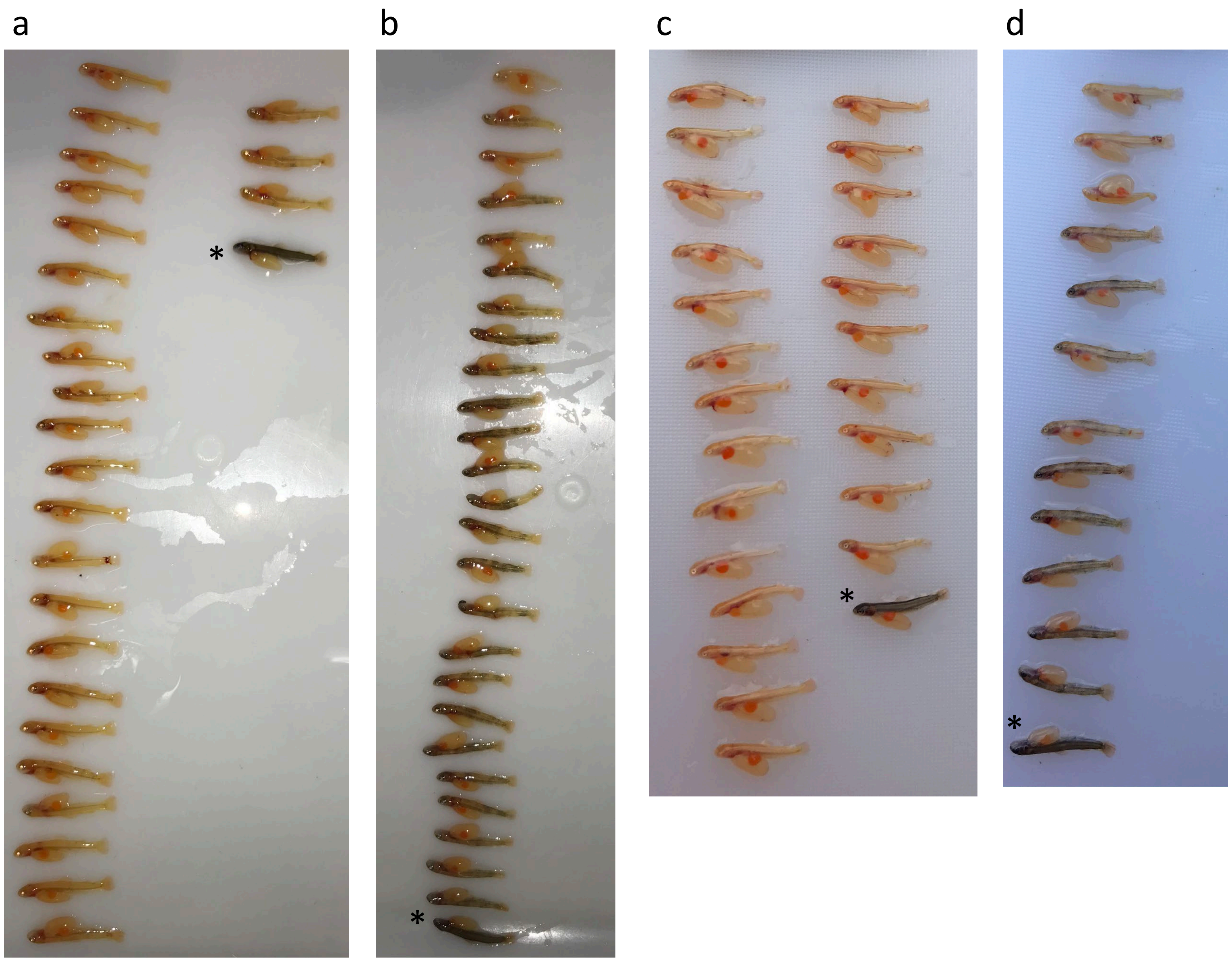


**Example of fry sampling based on visual inspection of pigmentation.**
**a** *slc45a2* FLAG KI (0.5 µM)
**b** *slc45a2* FLAG KI (4 µM)
**c** *dnd* SNR (0.15 µM)
**d** *dnd* SNR (4 µM)
*: wild type

### Supplementary Fig. 4.

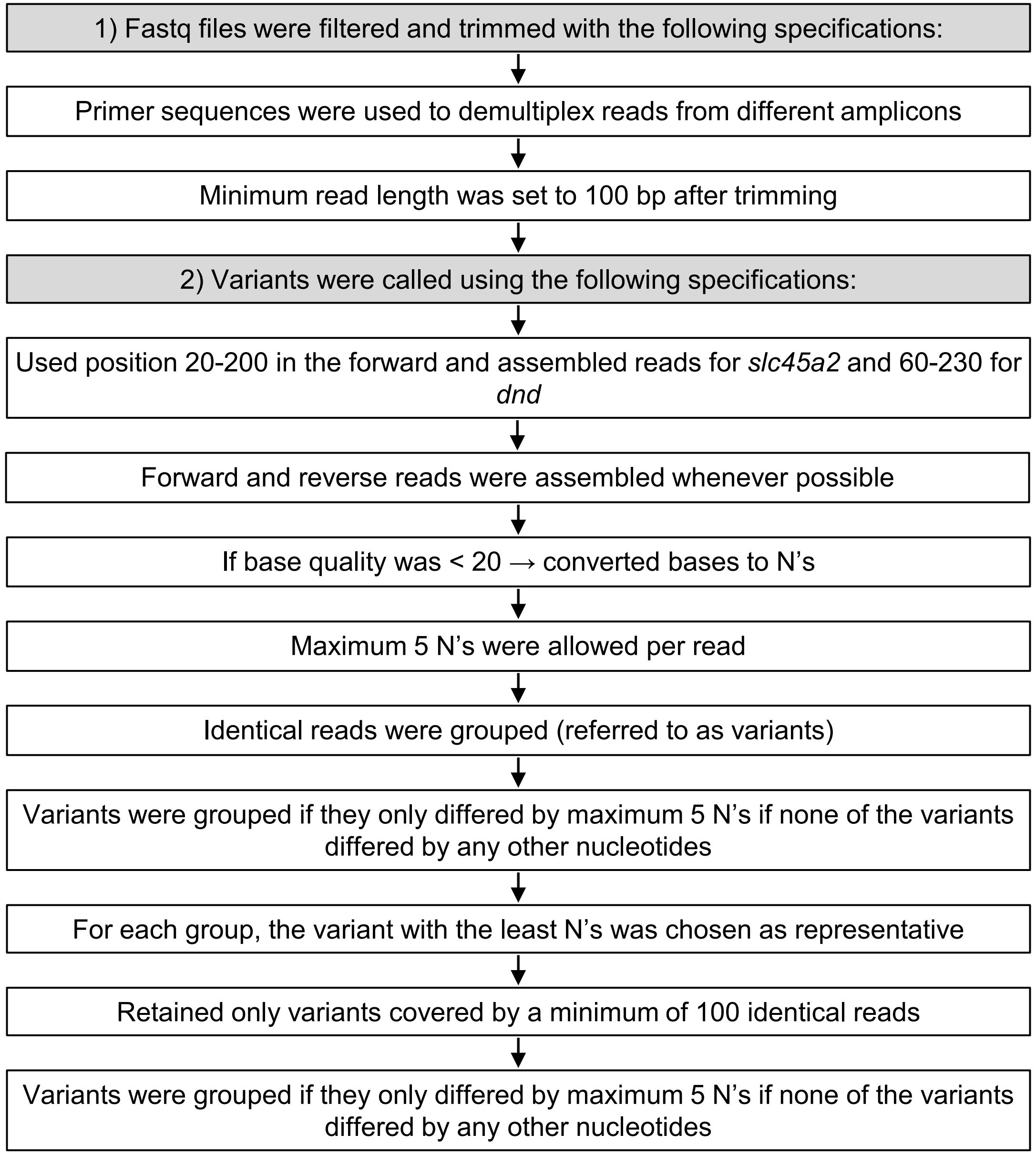


**Illustration of the settings applied for filtering, trimming and variant calling of the MiSeq reads.** Fastq files were filtered and trimmed with the following specifications; primer sequences were used to demultiplex reads from different amplicons on the same sequencing run, minimum read length was set to 100 bp, and forward and reverse reads were assembled to correct sequencing errors (minimum overlap between forward and reverse reads was set to 150 bp for *slc45a2* and 200 bp for *dnd* and allowing at most 20% mismatches between forward and reverse reads in the overlap region). Assembled reads were combined with forward reads that did not pass the assembly thresholds. Variants were then called using positions 20-200 for *slc45a2* and positions 60-230 for *dnd*. All bases with base quality < 20 were converted to N's, and maximum 5 N's were allowed per read. Identical reads were then grouped (referred to as variants) → variants that were only differing by up to 5 N's were grouped if none of the variants differed by any nucleotides → for each group the variant with the least N's was chosen as representative → only retained variants supported by a minimum of 100 reads → variants were grouped if they differed by up to 5 N's if none of the variants differed by any nucleotides. Finally, read counts were reported for the variants containing the inserted or edited sequence, separating those with a perfect match to the entire target sequence, and those with a correct insert sequence, but mismatches in the rest of the target sequence. In addition, read counts were reported wild type sequences.
