## Supplementary Table for "A refinement to gene editing in Atlantic salmon using asymmetrical oligonucleotide donors"

| **ODN** | **ODN concentration** | **Perfect HDR (%)** | **Erroneous HDR (%)** | **Other edits (%)** | **Wild type (%)** |
| --- | --- | --- | --- | --- | --- |
| *slc45a2*_FLAG_asym | 1.5 µM | 17.25 | 18.21 | 64.54 | 0 |
| *slc45a2*_FLAG_asym | 1.5 µM | 0.99 | 0 | 98.15 | 0.86 |
| *slc45a2*_FLAG_asym | 1.5 µM | 1.67 | 4.05 | 92.4 | 1.88 |
| *slc45a2*_FLAG_asym | 1.5 µM | 18.24 | 0.75 | 73.69 | 7.32 |
| *slc45a2*_FLAG_asym | 1.5 µM | 8.12 | 3.66 | 83.86 | 4.36 |
| *slc45a2*_FLAG_asym | 1.5 µM | 6.61 | 5.18 | 85.05 | 3.16 |
| *slc45a2*_FLAG_asym | 1.5 µM | 15.02 | 0 | 84.98 | 0 |
| *slc45a2*_FLAG_asym | 1.5 µM | 29.73 | 0 | 69.65 | 0.62 |
| *slc45a2*_FLAG_asym | 1.5 µM | 11.45 | 0 | 87.39 | 1.16 |
| *slc45a2*_FLAG_asym | 1.5 µM | 20.19 | 4.54 | 74.85 | 0.42 |
| *slc45a2*_FLAG_asym | 1.5 µM | 5.84 | 1.68 | 92.48 | 0 |
| *slc45a2*_FLAG_asym | 1.5 µM | 13.13 | 0.96 | 84.21 | 1.7 |
| *slc45a2*_FLAG_asym | 1.5 µM | 6.71 | 2.47 | 87.6 | 3.22 |
| *slc45a2*_FLAG_asym | 1.5 µM | **36.68** | 0.39 | 62.52 | 0.41 |
| *slc45a2*_FLAG_asym | 1.5 µM | 23.77 | 0 | 74.59 | 1.64 |
| *slc45a2*_FLAG_asym | 1.5 µM | 29.07 | 2.21 | 68.72 | 0 |
| *slc45a2*_FLAG_asym | 1.5 µM | 14.71 | 1.93 | 82.36 | 1 |
| *slc45a2*_FLAG_asym | 1.5 µM | 19.98 | 6.8 | 73.22 | 0 |
| *slc45a2*_FLAG_asym | 1.5 µM | 0.22 | 0.17 | 99.61 | 0 |
| *slc45a2*_FLAG_asym | 1.5 µM | 4.82 | 2.28 | 91.68 | 1.22 |
| *slc45a2*_FLAG_asym | 1.5 µM | 6.67 | 0.22 | 91.94 | 1.17 |
| *slc45a2*_FLAG_asym | 1.5 µM | 7.95 | 0 | 92.05 | 0 |
| *slc45a2*_FLAG_asym | 1.5 µM | 0.97 | 0 | 97.09 | 1.94 |
| *slc45a2*_FLAG_asym | 1.5 µM | **35.44** | 2.17 | 62.12 | 0.27 |
| *slc45a2*_FLAG_asym | 1.5 µM | 5.1 | 0.41 | 90.08 | 4.41 |
| *slc45a2*_FLAG_asym | 1.5 µM | 7.32 | 0 | 90.13 | 2.55 |
| *slc45a2*_FLAG_asym | 1.5 µM | 6.85 | 0.24 | 86.98 | 5.93 |
| *slc45a2*_FLAG_asym | 1.5 µM | 9.45 | 0.23 | 89.23 | 1.09 |
| *slc45a2*_FLAG_asym | 1.5 µM | **35.85** | 1.17 | 62.98 | 0 |
| *slc45a2*_FLAG_asym | 1.5 µM | 7.5 | 0.91 | 91.22 | 0.37 |
| Average |  | 13.58 | 2.02 | 82.85 | 1.56 |
| Std |  | 10.88 | 3.54 | 11.28 | 1.88 |
| *dnd1*_FLAG_asym | 1.5 µM | 0 | 0 | 100 | 0 |
| *dnd1*_FLAG_asym | 1.5 µM | **28.91** | 10.47 | 60.62 | 0 |
| *dnd1*_FLAG_asym | 1.5 µM | 5.54 | 0 | 94.46 | 0 |
| *dnd1*_FLAG_asym | 1.5 µM | 11.22 | 11.1 | 77.68 | 0 |
| *dnd1*_FLAG_asym | 1.5 µM | 0.89 | 0.29 | 98.82 | 0 |
| *dnd1*_FLAG_asym | 1.5 µM | 8.41 | 0 | 91.59 | 0 |
| *dnd1*_FLAG_asym | 1.5 µM | 0 | 0 | 100 | 0 |
| *dnd1*_FLAG_asym | 1.5 µM | 6.2 | 0 | 93.8 | 0 |
| *dnd1*_FLAG_asym | 1.5 µM | 5.91 | 76.28 | 17.81 | 0 |
| *dnd1*_FLAG_asym | 1.5 µM | 5.11 | 0 | 94.56 | 0.33 |
| *dnd1*_FLAG_asym | 1.5 µM | 0 | 0 | 100 | 0 |
| *dnd1*_FLAG_asym | 1.5 µM | 0 | 0 | 100 | 0 |
| *dnd1*_FLAG_asym | 1.5 µM | 0.4 | 8.08 | 91.52 | 0 |
| *dnd1*_FLAG_asym | 1.5 µM | 8.97 | 1.81 | 89.22 | 0 |
| *dnd1*_FLAG_asym | 1.5 µM | 1.71 | 9.29 | 89 | 0 |
| *dnd1*_FLAG_asym | 1.5 µM | 0 | 0 | 100 | 0 |
| *dnd1*_FLAG_asym | 1.5 µM | **32.73** | 0.78 | 66.49 | 0 |
| *dnd1*_FLAG_asym | 1.5 µM | 1.98 | 6.58 | 91.44 | 0 |
| *dnd1*_FLAG_asym | 1.5 µM | **29.08** | 1.64 | 69.28 | 0 |
| *dnd1*_FLAG_asym | 1.5 µM | 6.79 | 8.28 | 84.93 | 0 |
| *dnd1*_FLAG_asym | 1.5 µM | 3.87 | 5.6 | 90.53 | 0 |
| *dnd1*_FLAG_asym | 1.5 µM | 1.61 | 0.83 | 97.56 | 0 |
| *dnd1*_FLAG_asym | 1.5 µM | 0.84 | 0.83 | 98.33 | 0 |
| *dnd1*_FLAG_asym | 1.5 µM | 21.58 | 2.33 | 76.09 | 0 |
| Average |  | 7.57 | 6.01 | 86.41 | 0.01 |
| Std |  | 10.05 | 15.45 | 18.52 | 0.07 |
| *slc45a2*_FLAG_asym | 0.5 µM | 24.22 | 0 | 75.78 | 0 |
| *slc45a2*_FLAG_asym | 0.5 µM | 12.54 | 1.65 | 85.81 | 0 |
| *slc45a2*_FLAG_asym | 0.5 µM | 1.09 | 2.03 | 96.88 | 0 |
| *slc45a2*_FLAG_asym | 0.5 µM | 0 | 0 | 100 | 0 |
| *slc45a2*_FLAG_asym | 0.5 µM | 4.06 | 2.6 | 91.83 | 1.51 |
| *slc45a2*_FLAG_asym | 0.5 µM | 3.17 | 2.21 | 94.41 | 0.21 |
| *slc45a2*_FLAG_asym | 0.5 µM | 11.97 | 7.49 | 80.54 | 0 |
| *slc45a2*_FLAG_asym | 0.5 µM | 0.5 | 0 | 99.5 | 0 |
| *slc45a2*_FLAG_asym | 0.5 µM | 1.19 | 1.08 | 96.17 | 1.56 |
| *slc45a2*_FLAG_asym | 0.5 µM | 1.18 | 0 | 97.24 | 1.58 |
| *slc45a2*_FLAG_asym | 0.5 µM | 2.51 | 0 | 96.94 | 0.55 |
| *slc45a2*_FLAG_asym | 0.5 µM | 3.24 | 9.79 | 86.97 | 0 |
| *slc45a2*_FLAG_asym | 0.5 µM | 0 | 0 | 100 | 0 |
| *slc45a2*_FLAG_asym | 0.5 µM | 18.74 | 0.24 | 81.02 | 0 |
| *slc45a2*_FLAG_asym | 0.5 µM | 6.56 | 4.91 | 87.06 | 1.47 |
| *slc45a2*_FLAG_asym | 0.5 µM | 3.2 | 0.18 | 94.14 | 2.48 |
| *slc45a2*_FLAG_asym | 0.5 µM | 0.68 | 0.85 | 98.47 | 0 |
| *slc45a2*_FLAG_asym | 0.5 µM | 0 | 0 | 98.72 | 1.28 |
| *slc45a2*_FLAG_asym | 0.5 µM | 9.17 | 0 | 89.75 | 1.08 |
| *slc45a2*_FLAG_asym | 0.5 µM | 0.46 | 10.24 | 88.91 | 0.39 |
| *slc45a2*_FLAG_asym | 0.5 µM | 1.01 | 0 | 96.47 | 2.52 |
| *slc45a2*_FLAG_asym | 0.5 µM | 4.59 | 5.65 | 88.3 | 1.46 |
| *slc45a2*_FLAG_asym | 0.5 µM | 14.23 | 0 | 83 | 2.77 |
| Average |  | 5.40 | 2.13 | 91.65 | 0.82 |
| Std |  | 13.33 | 6.46 | 14.17 | 1.88 |
| *slc45a2*_FLAG_asym | 4.0 µM | 8.43 | 0 | 91.57 | 0 |
| *slc45a2*_FLAG_asym | 4.0 µM | 7.23 | 1.39 | 83.61 | 7.77 |
| *slc45a2*_FLAG_asym | 4.0 µM | 1.69 | 0.28 | 93.08 | 4.95 |
| *slc45a2*_FLAG_asym | 4.0 µM | 4.09 | 0.18 | 82.65 | 13.08 |
| *slc45a2*_FLAG_asym | 4.0 µM | 1.98 | 0 | 84.78 | 13.24 |
| *slc45a2*_FLAG_asym | 4.0 µM | 1.01 | 0.93 | 88.23 | 9.83 |
| *slc45a2*_FLAG_asym | 4.0 µM | 3.72 | 0 | 87.06 | 9.22 |
| *slc45a2*_FLAG_asym | 4.0 µM | 26.1 | 0 | 52.36 | 21.54 |
| *slc45a2*_FLAG_asym | 4.0 µM | 1.72 | 2.61 | 73.65 | 22.02 |
| *slc45a2*_FLAG_asym | 4.0 µM | 2.42 | 0.18 | 76.4 | 21 |
| *slc45a2*_FLAG_asym | 4.0 µM | 0.33 | 0.57 | 67.85 | 31.25 |
| *slc45a2*_FLAG_asym | 4.0 µM | 8.73 | 1.34 | 78.04 | 11.89 |
| *slc45a2*_FLAG_asym | 4.0 µM | 3.31 | 0 | 69.56 | 27.13 |
| *slc45a2*_FLAG_asym | 4.0 µM | 3.61 | 0.48 | 79.06 | 16.85 |
| *slc45a2*_FLAG_asym | 4.0 µM | 25.27 | 0 | 43.09 | 31.64 |
| *slc45a2*_FLAG_asym | 4.0 µM | 1.28 | 0 | 76.25 | 22.47 |
| *slc45a2*_FLAG_asym | 4.0 µM | 6.59 | 0.68 | 63.03 | 29.7 |
| *slc45a2*_FLAG_asym | 4.0 µM | 2.57 | 1.16 | 80.75 | 15.52 |
| *slc45a2*_FLAG_asym | 4.0 µM | 1.68 | 0.24 | 71.59 | 26.49 |
| *slc45a2*_FLAG_asym | 4.0 µM | 2.13 | 0 | 63.83 | 34.04 |
| *slc45a2*_FLAG_asym | 4.0 µM | 9.19 | 0.3 | 64.95 | 25.56 |
| *slc45a2*_FLAG_asym | 4.0 µM | 4.18 | 3.29 | 55.6 | 36.93 |
| *slc45a2*_FLAG_asym | 4.0 µM | 3.59 | 0.27 | 72.62 | 23.52 |
| Average |  | 5.69 | 0.60 | 73.90 | 19.81 |
| Std |  | 6.81 | 0.87 | 12.72 | 9.94 |
| *dnd1*_SE_asym | 1.5 µM | 0.84 | 0 | 99.16 | 0 |
| *dnd1*_SE_asym | 1.5 µM | 1.29 | 0 | 60.32 | 38.39 |
| *dnd1*_SE_asym | 1.5 µM | 0 | 0 | 100 | 0 |
| *dnd1*_SE_asym | 1.5 µM | 4 | 0 | 96 | 0 |
| *dnd1*_SE_asym | 1.5 µM | 6.06 | 3.25 | 90.69 | 0 |
| *dnd1*_SE_asym | 1.5 µM | 4.19 | 1.6 | 84.44 | 9.77 |
| *dnd1*_SE_asym | 1.5 µM | 0 | 0 | 97.3 | 2.7 |
| *dnd1*_SE_asym | 1.5 µM | 10.35 | 12.52 | 76.83 | 0.3 |
| *dnd1*_SE_asym | 1.5 µM | 43.57 | 18.15 | 38.28 | 0 |
| *dnd1*_SE_asym | 1.5 µM | 12.84 | 0 | 87.16 | 0 |
| *dnd1*_SE_asym | 1.5 µM | 14.08 | 57.53 | 28.39 | 0 |
| *dnd1*_SE_asym | 1.5 µM | 0 | 0.7 | 99.3 | 0 |
| *dnd1*_SE_asym | 1.5 µM | 16.83 | 14.81 | 68.36 | 0 |
| *dnd1*_SE_asym | 1.5 µM | 6.74 | 0 | 93.26 | 0 |
| *dnd1*_SE_asym | 1.5 µM | 18.43 | 2.56 | 79.01 | 0 |
| *dnd1*_SE_asym | 1.5 µM | 59.19 | 1.93 | 38.88 | 0 |
| *dnd1*_SE_asym | 1.5 µM | 8.15 | 21.38 | 70.47 | 0 |
| *dnd1*_SE_asym | 1.5 µM | 16.01 | 0 | 83.99 | 0 |
| *dnd1*_SE_asym | 1.5 µM | 29.02 | 6.24 | 64.74 | 0 |
| *dnd1*_SE_asym | 1.5 µM | 2.17 | 1.53 | 95.44 | 0.86 |
| *dnd1*_SE_asym | 1.5 µM | 6.22 | 30.46 | 63.32 | 0 |
| *dnd1*_SE_asym | 1.5 µM | 6.96 | 0 | 93.04 | 0 |
| *dnd1*_SE_asym | 1.5 µM | 2.19 | 0 | 96.83 | 0.98 |
| *dnd1*_SE_asym | 1.5 µM | 12.85 | 9.95 | 77.2 | 0 |
| *dnd1*_SE_asym | 1.5 µM | 2.73 | 0.37 | 90.74 | 6.16 |
| *dnd1*_SE_asym | 1.5 µM | 4.21 | 7.11 | 88.68 | 0 |
| Average |  | 12.47 | 8.26 | 78.36 | 0.90 |
| Std |  | 14.28 | 13.56 | 20.20 | 2.37 |
| *dnd1*_SE_asym | 0.15 µM | 0 | 0 | 100 | 0 |
| *dnd1*_SE_asym | 0.15 µM | 49.09 | 0 | 50.91 | 0 |
| *dnd1*_SE_asym | 0.15 µM | 17 | 0 | 83 | 0 |
| *dnd1*_SE_asym | 0.15 µM | 0 | 0 | 100 | 0 |
| *dnd1*_SE_asym | 0.15 µM | 47.35 | 18.06 | 34.59 | 0 |
| *dnd1*_SE_asym | 0.15 µM | 0 | 1.86 | 98.14 | 0 |
| *dnd1*_SE_asym | 0.15 µM | 9.67 | 0 | 89.94 | 0.39 |
| *dnd1*_SE_asym | 0.15 µM | 5.58 | 0 | 94.42 | 0 |
| *dnd1*_SE_asym | 0.15 µM | 0 | 0 | 100 | 0 |
| *dnd1*_SE_asym | 0.15 µM | 0.47 | 26.7 | 72.83 | 0 |
| *dnd1*_SE_asym | 0.15 µM | 0 | 0 | 100 | 0 |
| *dnd1*_SE_asym | 0.15 µM | 0 | 0 | 100 | 0 |
| *dnd1*_SE_asym | 0.15 µM | 0 | 0 | 100 | 0 |
| *dnd1*_SE_asym | 0.15 µM | 0 | 0 | 100 | 0 |
| *dnd1*_SE_asym | 0.15 µM | 0 | 0 | 100 | 0 |
| *dnd1*_SE_asym | 0.15 µM | 0 | 0 | 100 | 0 |
| *dnd1*_SE_asym | 0.15 µM | 0 | 0 | 100 | 0 |
| *dnd1*_SE_asym | 0.15 µM | 8.93 | 0 | 91.07 | 0 |
| *dnd1*_SE_asym | 0.15 µM | 0 | 1.66 | 98.34 | 0 |
| *dnd1*_SE_asym | 0.15 µM | 0 | 0 | 100 | 0 |
| *dnd1*_SE_asym | 0.15 µM | 0 | 5.14 | 94.86 | 0 |
| *dnd1*_SE_asym | 0.15 µM | 0 | 0 | 100 | 0 |
| *dnd1*_SE_asym | 0.15 µM | 1.92 | 1.97 | 96.11 | 0 |
| *dnd1*_SE_asym | 0.15 µM | 29.98 | 0 | 70.02 | 0 |
| Average |  | 7.39 | 2.41 | 90.18 | 0.02 |
| Std |  | 14.76 | 6.54 | 17.40 | 0.08 |
| *dnd1*_SE_asym | 4.0 µM | 2.12 | 9.86 | 88.02 | 0 |
| *dnd1*_SE_asym | 4.0 µM | 28.3 | 0 | 71.7 | 0 |
| *dnd1*_SE_asym | 4.0 µM | 0 | 0 | 100 | 0 |
| *dnd1*_SE_asym | 4.0 µM | 4.98 | 1.6 | 93.42 | 0 |
| *dnd1*_SE_asym | 4.0 µM | 9.32 | 0.39 | 88.93 | 1.36 |
| *dnd1*_SE_asym | 4.0 µM | 1.21 | 0.76 | 98.03 | 0 |
| *dnd1*_SE_asym | 4.0 µM | 7.6 | 21.36 | 71.04 | 0 |
| *dnd1*_SE_asym | 4.0 µM | 2.45 | 20.18 | 77.02 | 0.35 |
| *dnd1*_SE_asym | 4.0 µM | 5.98 | 1.19 | 92.83 | 0 |
| *dnd1*_SE_asym | 4.0 µM | 3.48 | 1.34 | 95.18 | 0 |
| *dnd1*_SE_asym | 4.0 µM | 25.65 | 0 | 74.35 | 0 |
| *dnd1*_SE_asym | 4.0 µM | 15.26 | 0.88 | 80.96 | 2.9 |
| Average |  | 7.36 | 3.27 | 86.50 | 0.20 |
| Std |  | 9.36 | 6.04 | 18.25 | 0.65 |
